# Nitrate activates a MKK3-dependent MAPK module via NLP transcription factors in Arabidopsis

**DOI:** 10.1101/2024.07.09.602786

**Authors:** Sebastian T. Schenk, Virginie Brehaut, Camille Chardin, Marie Boudsocq, Anne Marmagne, Jean Colcombet, Anne Krapp

## Abstract

Plant responses to nutrient availability are critical for plant development and yield. Nitrate, the major form of nitrogen in most soils, serves as both a nutrient and signaling molecule. Nitrate itself triggers rapid, major changes in gene expression, especially via NIN-LIKE PROTEIN (NLP) transcription factors, and stimulates protein phosphorylation. Mitogen-activated protein (MAP) kinase genes are among the early nitrate-responsive genes; however, little is known about their roles in nitrate signaling pathways. Here, we show that nitrate resupply to nitrogen-depleted Arabidopsis (*Arabidopsis thaliana*) plants triggers, within minutes, a MAPK cascade that requires NLP-dependent transcriptional induction of *MITOGEN-ACTIVATED PROTEIN KINASE KINASE KINASE 13* (*MAP3K13*) and *MAP3K14*. Importantly, nitrate reductase-deficient mutants exhibited nitrate-induced MAPK activities comparable to those observed in wild-type plants, indicating that nitrate itself is the signal that stimulates the cascade. We show that the modified expression of *MAP3K13* and *MAP3K14* affects nitrate-stimulated gene expression and modulates plant responses to nitrogen availability, such as nitrate uptake and senescence. Our finding that a MAPK cascade involving MAP3K13 and MAP3K14 functions in the complex regulatory network governing responses to nitrate availability will guide future strategies to optimize plant responses to nitrogen fertilization and nitrogen use efficiency.

**Significance statement:** Nitrate is an essential nutrient that also acts as a signaling molecule to regulate plant metabolism and development. We identified a specific MAPK cascade that is activated by nitrate and regulates several nitrate-dependent responses, such as senescence and nitrate transport.

## INTRODUCTION

Nitrogen (N) is a major macronutrient essential for all living organisms. Fungi and plants have the capacity to take up inorganic N from their environment and to metabolize it into organic N molecules. In addition to its important role as a plant nutrient, nitrate, the major form of N in most soils, also acts as a signal involved in many physiological and developmental pathways, such as leaf expansion (Walch-Liu et al., 2000), lateral root growth (Zhang and Forde, 2000), and seed dormancy (Alboresi et al., 2005). Understanding how plants perceive nitrate and how this perception is transduced into responses that optimize growth is important for the rational improvement of crop productivity and for mitigating pollution from the use of fertilizers.

Our understanding of the signaling network in response to nitrate availability has increased immensely in recent decades. The so-called primary nitrate response (PNR) is the rapid modification of the expression of roughly 2,000 genes in Arabidopsis (*Arabidopsis thaliana*) (Medici and Krouk, 2014). This gene regulatory network is orchestrated by several transcription factors (Vidal et al., 2020). Among these, members of the NODULE INCEPTION (NIN)-like protein family were identified as higher-level regulators in several species such as Arabidopsis and rice (*Oryza sativa*) (Castaings et al. 2009, Konishi and Yanagisawa, 2013; Marchive et al. 2013; Alfatih et al., 2020; Wu et al., 2021; Zhang et al., 2022). In Arabidopsis, NIN-LIKE PROTEIN 7 (NLP7) and NLP2 are the two members of this family that contribute most to the regulation of the PNR in seedlings and adult plants (Castaings et al., 2009; Konishi et al., 2021; Durand et al., 2023). NLP6, the closest homolog of NLP7, can partially compensate for the loss of NLP7 function in the single *nlp7* mutant. Nonetheless, *nlp6* single mutants show only minor modifications of traits related to N availability (Cheng et al., 2023). Interestingly, NLP7, NLP6, and NLP2 are regulated by nuclear retention as quickly as 3 min after nitrate resupply (Marchive et al., 2013; Durand et al., 2023; Cheng et al., 2023). NLP7 was recently shown to act as a nitrate sensor via the direct binding of nitrate to its N -terminal domain, which is required to activate transcriptional regulation (Liu et al., 2022). This nitrate-binding domain is conserved among all NLPs. Nitrate is also sensed by the transceptor NRT1/PTR FAMILY 6.3 (NPF6.3, also reported as NITRATE TRANSPORTER 1.1 [NRT1.1] and CHLORINA 1 [CHL1]), a dual-affinity nitrate transporter that monitors changes in external nitrate levels (Ho et al., 2009) in plants grown in the presence of ammonium prior to nitrate addition (Wang et al., 2009).

Besides transcriptional regulation, early responses to nitrate resupply also involve post-translational modifications such as protein phosphorylation (Liu et al., 2020; Wang et al., 2021). Large changes in the phosphoproteome occur in response to N availability (Engelsberger and Schulze, 2012; Menz et al., 2016). Reversible protein phosphorylation can rapidly regulate the localization, stability, activity, and interaction partners of target proteins (Yip Delormel and Boudsocq, 2019). Studies performed over 25 years ago using inhibitors of protein phosphatases or kinases suggested that protein phosphorylation and dephosphorylation might be involved in nitrate signaling (Sakakibara et al., 1997; Sueyoshi et al., 1999). Since then, two CBL-INTERACTING PROTEIN KINASEs (CIPKs) have been shown to contribute to nitrate signaling by forming a module with Calcineurin B-Like 1 (CBL1) and CBL9. CIPK8 positively regulates the low-affinity nitrate response (Hu et al., 2009), while CIPK23 phosphorylates NPF6.3 under high-affinity nitrate conditions (Ho et al., 2009).

In addition, subgroup III calcium-dependent protein kinases (CPK10, CPK30, and CPK32) phosphorylate the key transcription factor NLP7 in response to a calcium-mediated nitrate signal. This phosphorylation at Ser-205, which is conserved among NLP transcription factors, leads to the nuclear retention of NLP7 (Liu et al., 2017). In agreement with this finding, pretreating seedlings with inhibitors of calcium signaling diminished nitrate-triggered changes in gene expression (Riveras et al., 2015, Liu et al., 2017; Adavi and Sathee, 2024). Importantly, not all nitrate-stimulated changes were abolished by these inhibitor treatments, suggesting that both calcium-dependent and -independent nitrate mechanisms co-exist for the regulation of gene expression.

Protein phosphorylation by mitogen-activated protein kinases (MAPK) is associated with plant development and responses to biotic and abiotic stress (Pitzschke et al., 2009; Zhang and Zhang, 2022). The minimal MAPK pathway consists of three core members: a MAPK (MPK), a MAPK kinase (MAP2K or MKK), and a MAP2K kinase (MAP3K). MAP3Ks phosphorylate and thereby activate MAP2Ks, which in turn phosphorylate and activate their downstream MAPKs. These MAPK cascades are conserved in eukaryotes including yeast (*Saccharomyces cerevisiae*), mammals, and plants, where they convert extracellular signals into cellular responses. Genes encoding 20 MAPKs, 10 MAP2Ks, and ∼80 putative MAP3Ks belonging to three distinct families have been identified in the Arabidopsis genome based on their similarity to their animal and fungal counterparts. The MAPK and MAP2K families are both organized into four subclades (groups A–D). Among the three MAP3K families, only MAPK/extracellular signal-regulated kinase (ERK) kinase kinase (MEKK)-like kinases (21 members) are unambiguously involved in the activation of plant MAP2Ks (Xu and Zhang, 2015). These MEKK-like kinases are themselves organized into three subclades (Colcombet et al., 2016). In particular, genes of subclade III (*MAP3K13–21*) show increased transcript levels in response to abiotic and biotic factors, and their encoded proteins interact with MKK3 to activate group C MAPKs (Sözen et al., 2020; Danquah et al, 2015). However, functional studies of these MAP3Ks in plants are still scarce. Among the conclusions from such studies are that MAP3K17 and MAP3K18 function in an abscisic acid (ABA)-activated MAPK module during plant adaptation to drought (Danquah et al., 2015; Matsuoka et al., 2015) and that MAP3K14, MAP3K17, and MAP3K21 contribute to the plant defense response against herbivores (Hernandez and Martinez, 2022; Sözen et al., 2020). Recent data also suggest that these MAP3Ks play a role in modulating seed dormancy (Regnard et al., 2024; Otani et al., 2024; Chen et al., 2023), and overexpressing several of these genes altered the ABA sensitivity of seed germination as well as the stress responses of Arabidopsis seedlings (Choi et al., 2017). Few proteins have been proposed to be targeted by MKK3 modules downstream of group C MAPKs. *In vitro*, MPK1 phosphorylates ROP BINDING PROTEIN KINASE 1 (RBK1), and analysis of knock-out mutants suggested that a MPK1-MKK3-RBK1 module plays a role in auxin-induced cell expansion (Enders et al., 2017). In the context of seed dormancy, the MPK7-mediated phosphorylation of ETHYLENE RESPONSE FACTOR 4 (ERF4) leads to its degradation and the expression of EXPA proteins involved in radicle protrusion (Chen et al., 2024).

Our knowledge of MAPK cascades that participate in N-related signaling pathways is limited. Forde et al. (2013) showed that MEKK1 regulates root elongation in Arabidopsis in response to L-glutamate. Moreover, Arabidopsis MKK9 is thought to negatively regulate anthocyanin biosynthesis under N limitation–triggered senescence by promoting nitrate acquisition (Luo et al., 2017). Interestingly, the clade III MAP3K genes *MAP3K13* and *MAP3K14* are strongly upregulated during the PNR, and this regulation at least partially depends on NLP7. Indeed, NLP7 binds to the promoter regions of *MAP3K13* and *MAP3K14* (Marchive et al., 2013; Alvarez et al., 2020). Based on the previous finding that the transcriptional regulation of clade III *MAP3K*s is indicative of the activation of an MKK3 module (Colcombet et al., 2016), these data led us to hypothesize that nitrate may activate an MKK3 module in an MAP3K13/14- and NLP-dependent manner.

In the present work, we show that the activity of MPK1, MPK2, and MPK7 is stimulated by nitrate resupply in Arabidopsis and that the nitrate-triggered activation of MPK7 is dependent on NLP transcription factors, MAP3K13, MAP3K14, and MKK3, revealing a yet undescribed nitrate- and NLP2/7-dependent MAPK signaling cascade in seedlings. We also provide evidence that MAP3K13 and MAP3K14 regulate nitrate uptake and N deficiency– triggered senescence.

## RESULTS

### Nitrate resupply of N-depleted seedlings rapidly induces *MAP3K13* and *MAP3K14* expression

Nitrate resupply to N-depleted seedlings triggers a rapid change in the expression levels of hundreds of genes (Wang et al., 2001; Scheible et al., 2004; Orsel et al., 2005; Vidal et al., 2020). Among these, *MAP3K13* and *MAP3K14* expression is highly nitrate-responsive, and this transcriptional induction is under the direct control of the transcription factor NLP7 (Marchive et al., 2013; Alvarez et al., 2020). We hypothesized that a MAPK cascade initiated by MAP3K13 and/or MAP3P14 might become activated by nitrate resupply and participate in the regulation of nitrate-triggered responses. In agreement with this hypothesis, a signal-triggered increase in the transcript levels of clade-III *MAP3K*s has been reported as an obligatory step to activate MAP2K, MKK3, and the group C MAPKs MPK1, MPK2, MPK7, and MPK14 (Danquah et al., 2015; Sözen et al., 2020).

To characterize the kinetics of their transcript accumulation, we performed a time-course analysis of *MAP3K13* and *MAP3K14* expression in young Arabidopsis seedlings by reverse-transcription quantitative PCR (RT-qPCR) (Figure 1a). We grew seedings under replete N supply (3 mM nitrate) for 10 days, followed by 3 days of N starvation before adding either 3 mM potassium nitrate (KNO_3_) or potassium chloride (KCl) to the liquid growth medium, followed by sampling from 2.5 min to 45 min. *MAP3K13* and *MAP3K14* were expressed at very low levels under N-starvation conditions and after KCl treatment. Nitrate resupply led to a rapid and transient increase in *MAP3K13* and *MAP3K14* expression, with a significant increase after 5 min and a peak corresponding to an 18-fold and 32-fold induction, respectively, at 10–15 min. Expression dropped close to initial levels at 45 min after nitrate resupply (Figure 1b). The transcript levels of other clade-III *MAP3K* genes did not strongly increase in response to nitrate resupply (Figure S1). For comparison, the transcript levels of genes encoding enzymes involved in nitrate assimilation, such as *NITRATE TRANSPORTER 2.1 (NRT2.1)* and *NITRITE REDUCTASE* (*NIR*), increased less rapidly, reaching peak levels at 30 min after nitrate resupply and remaining high thereafter (Figure 1c). The expression levels of genes encoding regulators of N responses, such as the transcription factor HYPERSENSITIVE TO LOW PI-ELICITED PRIMARY ROOT SHORTENING 1 (HRS1), showed a rapid response, with a transient increase, first detectable at 10 min and peaking at 15 min, before declining. Thus, the expression profiles of *MAP3K13* and *MAP3K14* in response to nitrate resupply suggest that their encoded MAP3Ks might play a role in the rapid nitrate signaling pathway.

**Figure 1.**
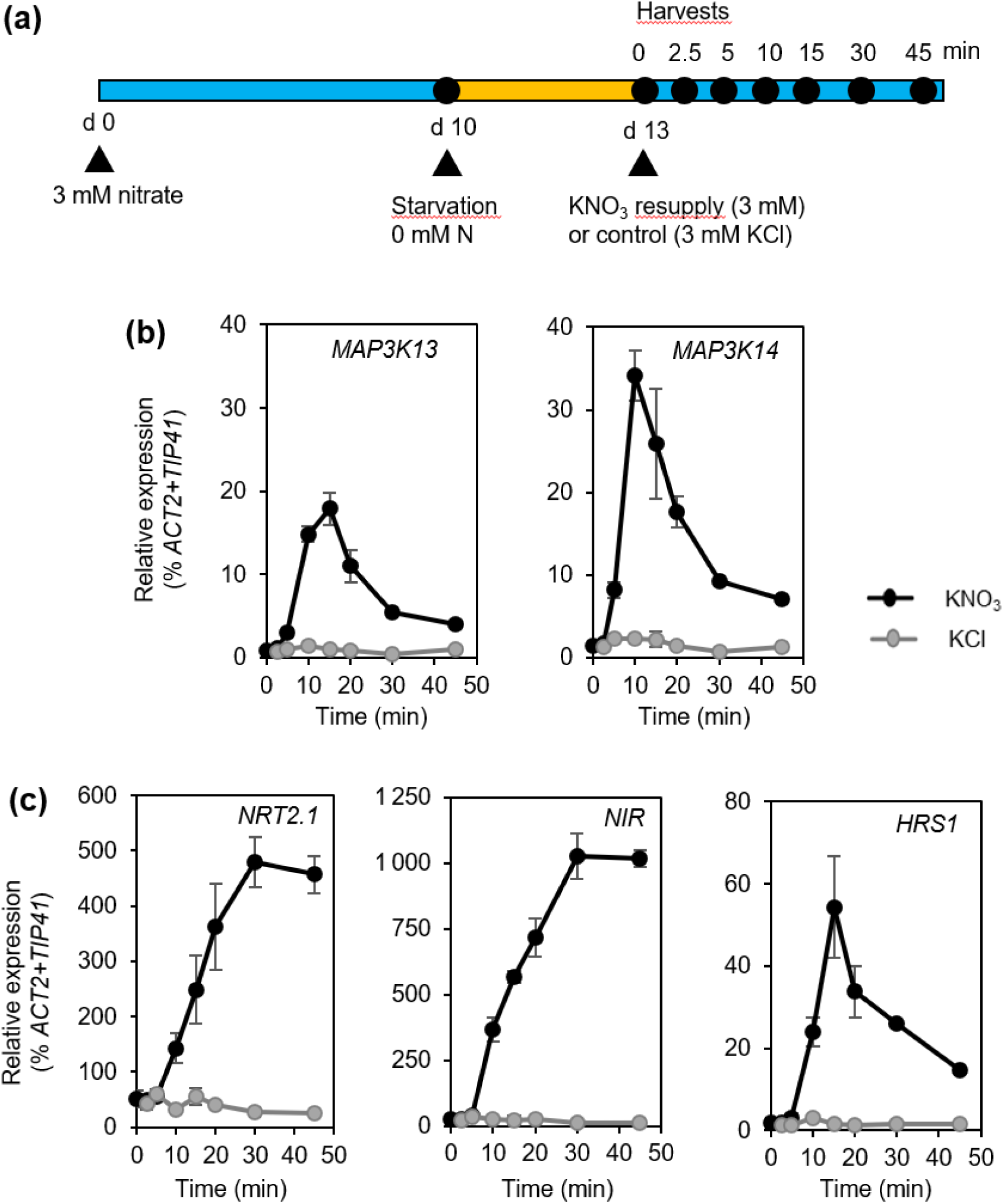
Transcript levels of *MAP3K13* and *MAP3K14* rapidly, strongly, and transiently increase after nitrate resupply to N-depleted seedlings. (a) Experimental design. (b, c) Relative transcript levels of the indicated genes in seedlings, as measured by RT-qPCR and normalized using *TIP41* (At4g34270) and *ACTIN2* (*ACT2*, At3g18780) as control genes. d = day. Data are means ± standard error of the mean (SEM, n = 3 independent samples of 30 seedlings each) for one representative experiment out of two.

### Nitrate resupply triggers the activation of group C MAPKs

Clade-III MAP3Ks, notably MAP3K13 to MAP3K20, were previously shown to interact with MKK3 in yeast two-hybrid (Y2H) assays (Sözen et al., 2020). They also activated the group C MAPK MPK2 in an MKK3-dependent manner when their encoding constructs were transiently expressed in Arabidopsis mesophyll protoplasts (Sözen et al., 2020). Furthermore, stimulation of *MAP3K18* and *MAP3K14* expression in plants led to the activation of some group C MAPKs in a strictly MKK3-dependent manner (Danquah et al., 2015; Sözen et al., 2020). We thus wondered if the nitrate-induced increases in *MAP3K* transcript levels in N-starved seedlings might trigger the activation of a MAPK cascade composed of MKK3 and group C MAPKs. Using specific anti-MAPK antibodies (Ortiz-Masia et al., 2007, Doczi et al., 2007), we immunoprecipitated three of the four group C MAPKs individually (MPK1, MPK2, and MPK7) from Arabidopsis seedlings (using wild-type [WT] Col-0) after nitrate resupply and assayed their activity as the capacity to phosphorylate myelin basic protein (MBP), a generic MAPK substrate. The activities of these three MAPKs increased starting at 10 min following the beginning of nitrate resupply, with a peak at 15–20 min (Figure 2a). Their activity did not increase after the addition of 3 mM KCl, pointing to the specificity of this response to nitrate addition.

**Figure 2.**
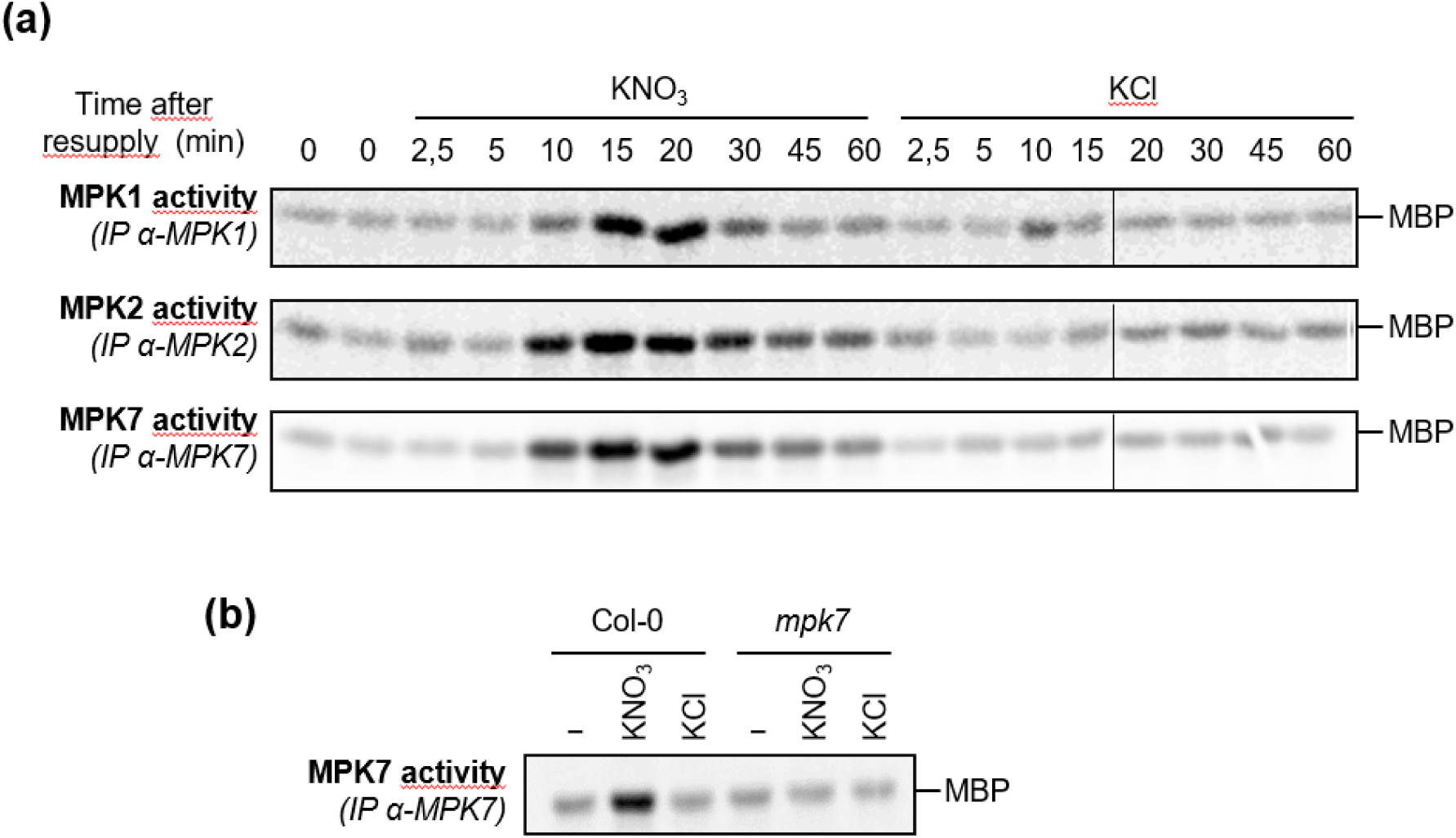
Activation of group C MAPKs by nitrate. (a) Kinase activity of MPK1, MPK2, and MPK7 after immunoprecipitation (IP) with specific antibodies from N-depleted Col-0 seedlings resupplied with 3 mM KNO_3_ or KCl for the indicated durations and following the experimental setup shown in Figure 1a. (b) Kinase activity of MPK7 after immunoprecipitation (IP) with specific antibodies from N-depleted Col-0 or *mpk7* seedlings before (−) and after resupply with 3 mM KNO_3_ or KCl for 15 min.

We selected MPK7 for further study and measured its activity as a proxy of the activation of the entire module. A T-DNA insertion mutant in *MPK7* showed impaired nitrate-induced immunoprecipitated kinase activity (Figure 2b). To rule out the possibility that the increased kinase activity in Col-0 seedlings was due to increased MPK7 protein abundance, we assessed MPK7 accumulation by immunoblotting. Since the anti-MPK7 antibody is unable to detect MPK7 in immunoblots (Doczi et al., 2007), we utilized a transgenic line expressing the *MPK7* locus cloned in-frame with the sequence encoding an HA (human influenza hemagglutinin) tag under the control of the *MPK7* promoter to monitor MPK7-HA abundance (Sözen et al., 2020). Immunoprecipitation with anti-HA antibodies confirmed a transient increase in MPK7-HA activity with similar kinetics to endogenous MPK7 in Col-0 (Figure S2). Importantly, MPK7-HA protein levels remained constant over the time course (Figure S2). Thus, we conclude that nitrate resupply triggers the activation of MPK1, MPK2, and MPK7 within minutes rather than an increase in their abundance.

### The activation of MPK7 is specific to nitrate and sensitive to micromolar nitrate levels

To further characterize nitrate-triggered MPK7 activation, we measured its sensitivity toward nitrate via a dose-response experiment using the *MPK7-HA* transgenic line. In agreement with earlier findings on nitrate-regulated gene expression (Wang et al., 2007), a nitrate concentration as low as 3 µM triggered the activation of MPK7-HA (Figure 3a). Next, we asked whether MPK7 activation was specific to nitrate as the sole N source. Indeed, neither ammonium nor glutamine, provided as 3 mM NH_4_Cl and 1.5 mM glutamine, respectively, activated MPK7-HA (Figure 3b). To confirm that the assimilation of nitrate into organic N-containing compounds was not required to activate this MAPK cascade, we examined MPK7 activation in the *nia1 nia2* mutant, which exhibits greatly decreased nitrate reductase activity (only 0.5% of WT). In this mutant that barely assimilates nitrate (Wilkinson and Crawford, 1993), the activation of endogenous MPK7 by nitrate was similar to that in WT (Figure 3c). Thus, the activation of MPK7 is specific and highly sensitive to nitrate and is triggered by nitrate itself.

**Figure 3.**
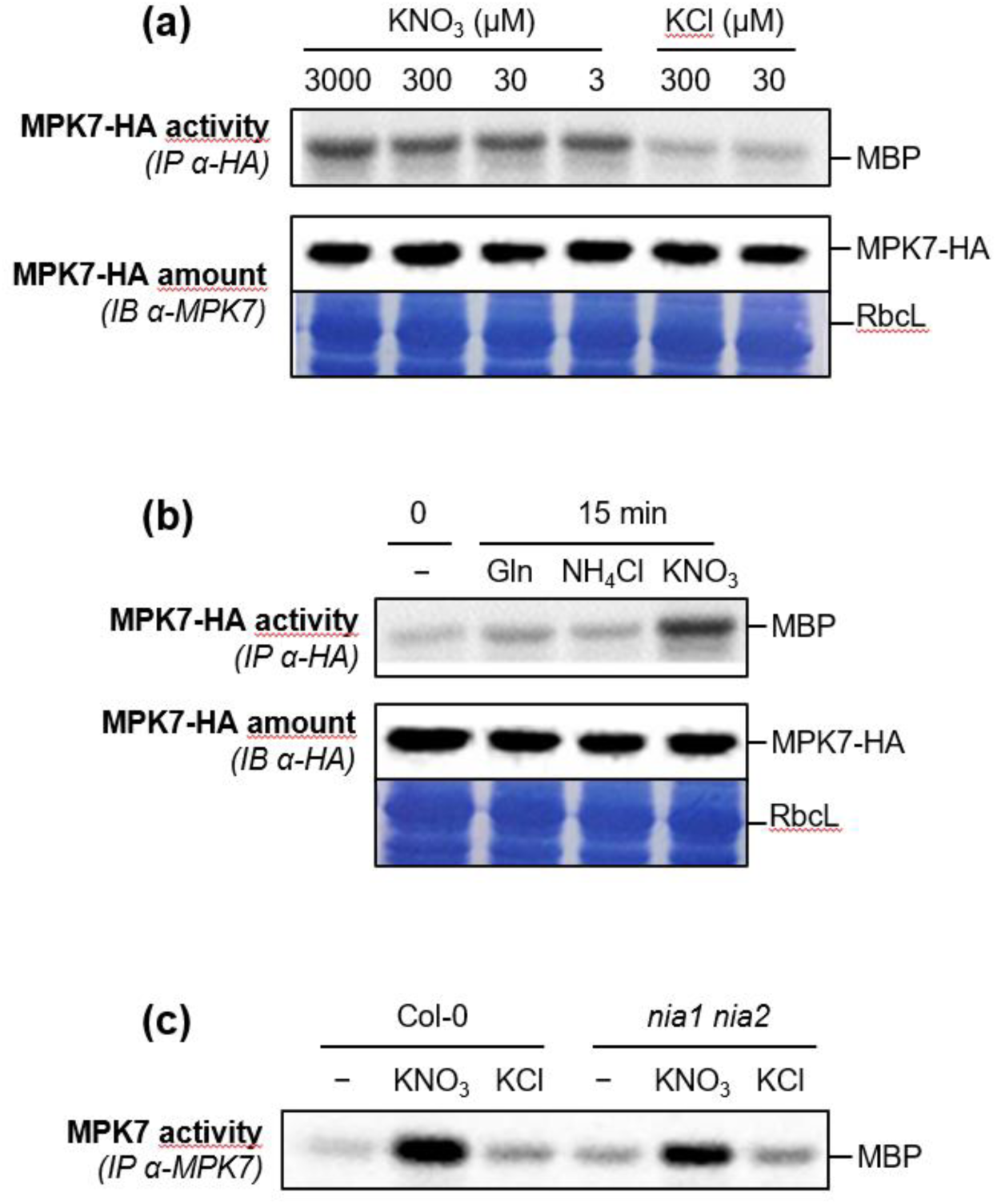
Nitrate-triggered MPK7 activation is sensitive to and specific for nitrate and independent of nitrate reductase activity. (a and b) Kinase activity of MPK7-HA after immunoprecipitation (IP) with an anti-HA antibody from N-depleted seedlings expressing *MPK7-HA* before (−) and after resupply with various µM concentrations of KNO_3_ or KCl as indicated (a) or 1.5 mM glutamine, 3 mM NH_4_Cl, or 3 mM KNO_3_ (b) for 15 min. Protein abundance was monitored by immunoblotting (IB) using an anti-HA antibody. Equal loading is indicated by Coomassie staining of the membrane. RbcL, Rubisco large subunit. (c) Kinase activity of MPK7 after immunoprecipitation (IP) with specific antibodies from N-depleted Col-0 and *nia1 nia2* seedlings before (−) and after resupply with 3 mM KNO_3_ or KCl for 15 min.

### Loss of NPF6.3 nitrate transport activity diminishes nitrate-triggered MPK7 activation

The nitrate transporter NPF6.3 (also reported as NRT1.1 and CHL1) is a nitrate sensor (Ho et al., 2009). To test whether the nitrate-triggered activation of MPK7 depends on the nitrate-sensing function of NPF6.3, we measured MPK7 activity in *chl1-5* and *chl1-9* mutants. Whereas the T-DNA insertion knockout mutant *chl1-5* lacks both nitrate transport and nitrate sensing, the *chl1-9* mutant is only impaired in nitrate transport. The function of NPF6.3 as a nitrate sensor in regulating the PNR depends on the N source supplied during growth before nitrate addition. Indeed, while NPF6.3 was shown to be dispensable for the response to nitrate of seedlings that experienced a complete 3-day N starvation, NPF6.3 is required for the PNR in seedlings cultivated in the presence of ammonium prior to nitrate supply (Wang et al., 2009). We thus examined nitrate-triggered MPK7 activation in seedlings after N starvation following the protocol described in Figure 1a, and separately in seedlings cultivated on medium containing 1.5 mM ammonium succinate for 13 days, comparing WT to the *chl1-5* and *chl1-9* mutants (Figure 4a, b). In N-depleted seedlings, we observed no difference between WT and either *chl1* mutant; by contrast, seedlings grown on ammonium-containing medium before nitrate addition exhibited a strongly decreased but not abolished stimulation of MPK7 activity in both *chl1* mutants compared to WT. Thus, nitrate transport, but not nitrate sensing, by NPF6.3 is at least partially required for the activation of this MAPK cascade in response to nitrate in seedlings that have not experienced a N-starvation treatment.

**Figure 4.**
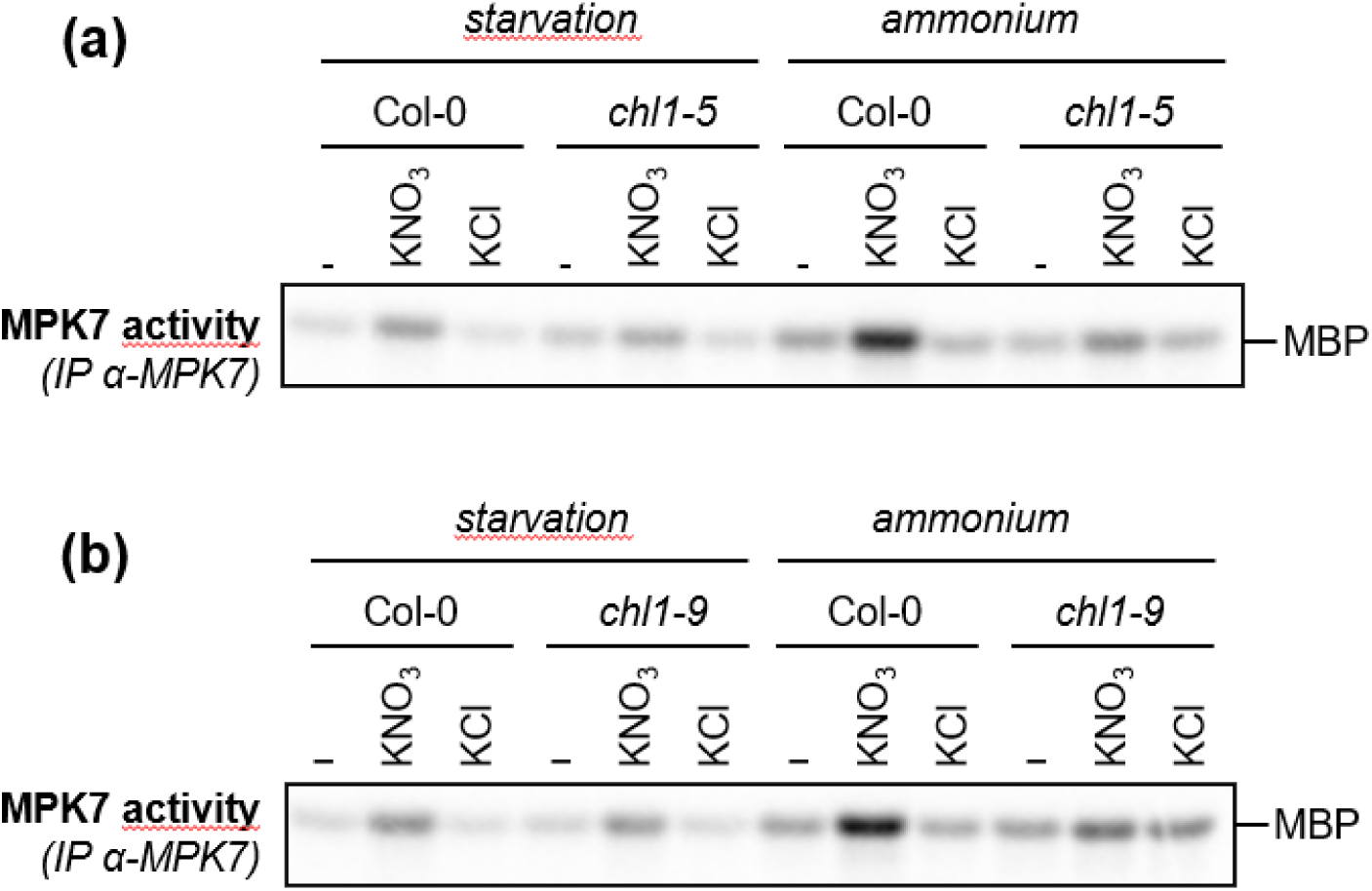
Loss of function of the nitrate transceptor NPF6.3 diminishes the nitrate-triggered induction of MPK7 activity in seedlings grown on ammonium succinate. (a and b) Kinase activity of MPK7 in the indicated genotypes after immunoprecipitation (IP) with specific antibodies from N-depleted seedlings (“starvation”) and seedlings grown on 1.5 mM ammonium succinate (“ammonium”) before (−) and after resupply with 3 mM KNO_3_ or KCl for 15 min.

### The activation of group C MAPKs by nitrate depends on MAP3K13, MAP3K14, and MKK3

We wanted to identify the upstream kinases of the nitrate-activated MAPK module. In fact, clade-III MAP3Ks and MKK3 are often involved in group C MAPK activation. We previously showed that this activation occurs due to the upregulation of *MAP3Ks* (Danquah et al., 2015; Sözen et al., 2020). Because *MAP3K13* and *MAP3K14* transcripts strongly accumulate upon nitrate treatment (Figure 1), we hypothesized that among the eight clade-III MAP3Ks, MAP3K13 and MAP3K14 play major roles in the activation of the module. Accordingly, we created two independent double mutant lines, referred to hereafter as *map3k13 map3k14-cr#1* (*map3k13^578insC^ map3k14^551insT^*) and *map3k13 map3k14-cr#2 (map3k13^578insT^ map3k14^551insA^)*, by CRISPR (clustered regularly interspaced short palindromic repeats)/Cas9 (CRISPR-associated protein 9)-mediated gene editing (Figure 5a). Both double mutants contain single-base insertions 578 bp downstream from the start codon of *MAP3K13* and 551 bp downstream from the start codon of *MAP4K14*, introducing frameshifts in the open reading frame and premature stop codons. Since at that time we did not have anti-MPK7 antibody remaining, we crossed the *map3K13 map3k14-cr1* mutant line to the *MPK7-HA* line to allow MPK7-HA immunoprecipitation in the double mutant background. The nitrate-induced activation of MPK7-HA was totally abolished in this background (Figure 5b). To test the relative role of each MAP3K, we identified their respective single mutants in a backcross of the *map3K13 map3k14-cr#1 MPK7-HA* line to Col-0. Gene-edited mutants of *MAP3K13* or *MAP3K14* showed an intermediate response between that of Col-0 and the double mutant, with diminished activation of MPK7-HA in response to nitrate (Figure 5c). We subsequently produced new specific anti-MPK7 antibodies and confirmed the suppression of nitrate-triggered MPK7 activation in the two *map3K13 map3k14-cr* double mutants based on immunoprecipitation of endogenous MPK7 (Figure S3). Nitrate-triggered MPK7 activation was also abolished in two *mkk3* mutants (Figure 5d) and restored in the *mkk3-1* mutant complemented by *MKK3-YELLOW FLUORESCENT PROTEIN (YFP)*.

**Figure 5.**
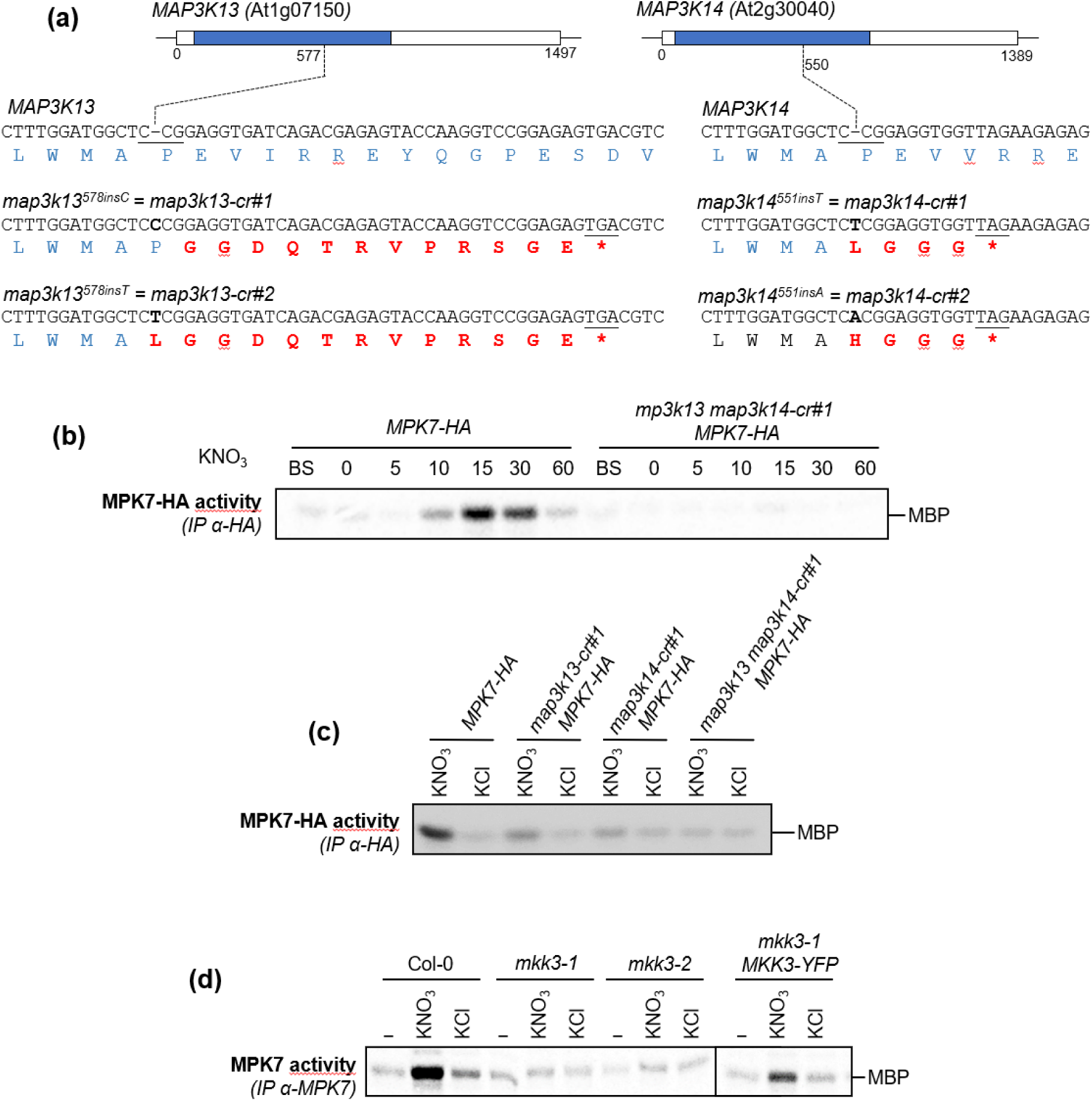
MAP3K13, MAP3K14, and MKK3 are required for the induction of MPK7 activity by nitrate. (a) Genomic structures of *MAP3K13* and *MAP3K14* and summary of the gene-edited lines used in this work. The sequence encoding the kinase domain is indicated in blue. (b and c) Kinase activity of MPK7-HA after immunoprecipitation (IP) with an anti-HA antibody from N-depleted seedlings expressing *MPK7-HA* in Col-0 or the *map3k13 map3k14-cr#1* mutant before starvation, after 3 days of N starvation (−), and after resupply with 3 mM KNO_3_ or KCl for the indicated time (b) or for 15 min (c). (d) Kinase activity of MPK7 after immunoprecipitation (IP) with specific antibodies from N-depleted seedlings of the indicated genotypes before (−) and after resupply with 3 mM KNO_3_ or KCl for 15 min.

To verify that a phosphorylation cascade is indeed the only molecular mechanism leading to the observed increase in MAPK activity, we measured the transcript levels of group C *MAPK*s and *MKK3* by RT-qPCR following nitrate resupply in a time course. We detected no major changes in their expression levels during the early nitrate response (Figure S4). Taken together, these results indicate that nitrate resupply to N-starved seedlings triggers a MAPK cascade within minutes in a manner strictly dependent on MAP3K13, MAP3K14, and MKK3.

### Nitrate-regulated *MAP3K13* and *MAP3K14* expression and the activation of the MAPK cascade depend on NLP-type transcription factors

To elucidate the roles of the transcription factors NLP2, NLP6, and NLP7, which function as master regulators of nitrate signaling, we measured *MAP3K13* and *MAP3K14* expression levels following nitrate resupply in single and double mutants of *NLP2*, *NLP6*, and *NLP7*. While *nlp6-1* showed almost no differences compared to WT, *nlp2-1* and *nlp7-1* both displayed a diminished transient induction of *MAP3K13* and *MAP3K14* expression, reaching 52% and 56%, respectively for *nlp2-1* and reaching 41% and 39%, respectively for *nlp7-1*, of the transcript levels seen in Col-0 at the expression peak (Figure 6a). In the *nlp6-2 nlp7-1* and *nlp2-1 nlp7-1* double mutants, *MAP3K13* and *MAP3K14* mRNA levels were slightly lower than those in the *nlp7-1* mutant in response to nitrate (Figure 6a). The effects on gene expression in the double mutants were only slightly stronger than those in the *nlp7* and *nlp2* single mutants, suggesting that NLP2 and NLP7 homodimers and/or heterodimers regulate *MAP3K13* and *MAP3K14* expression. These results are in line with the previous finding that NLP7 and/or NLP2 directly binds to the promoters of *MAP3K13* and *MAP3K14* (Marchive et al., 2013; Alvarez et al., 2020; Durand et al., 2023).

**Figure 6.**
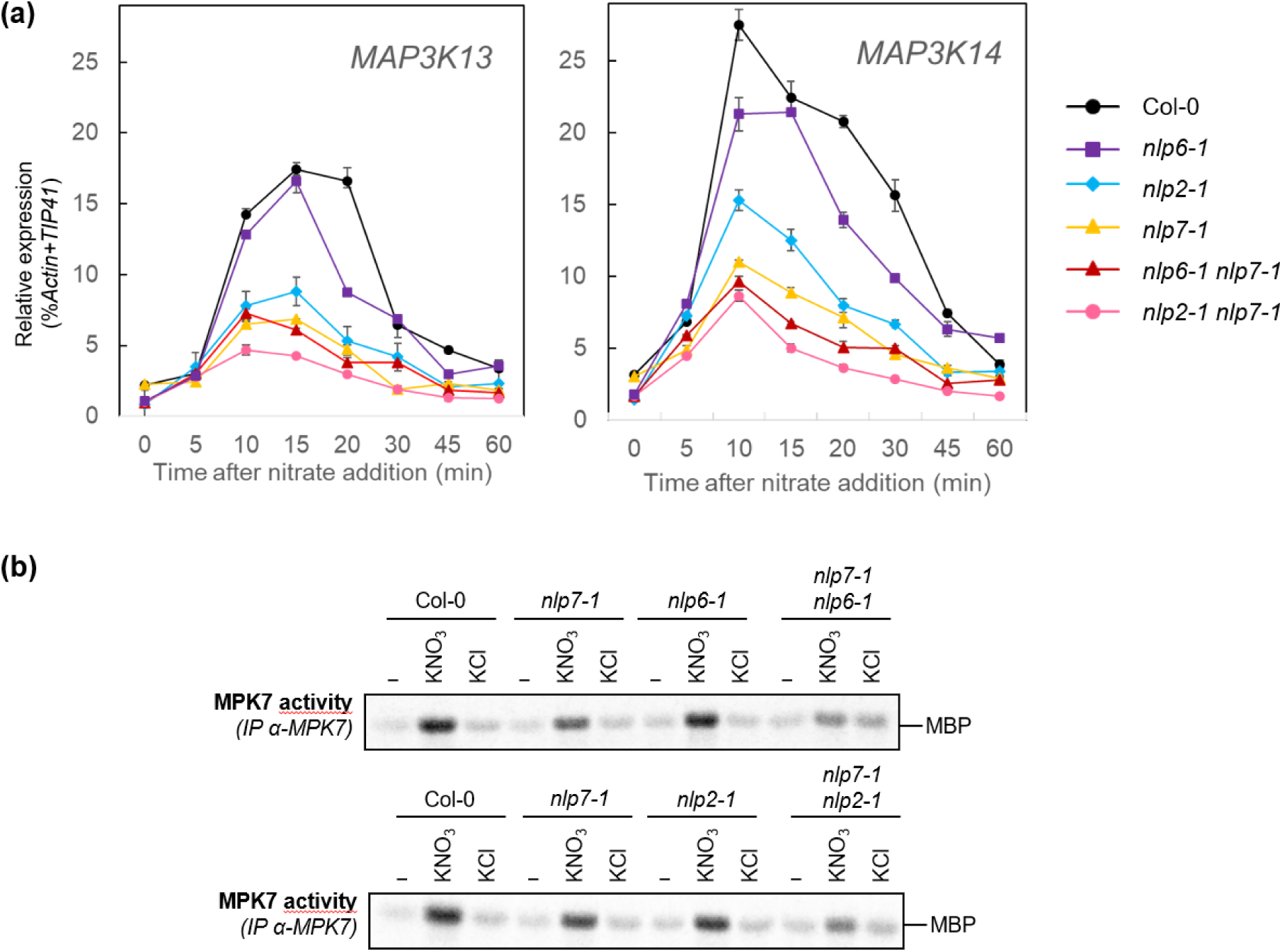
Rapid induction of *MAP3K13* and *MAP3K14* gene expression and the activation of MPK7 by nitrate depend on NLP transcription factors. (a) Relative transcript levels of *MAP3K13* and *MAP3K14* after nitrate resupply, as measured by RT-qPCR of Col-0 and *nlp* mutant seedlings grown in 6-well plates for 13 days (10 days on 3 mM KNO_3_ followed by 3 days on 0 mM KNO_3_). Seedlings were resupplied with 3 mM KNO_3_ or KCl for 5, 10, 15, 20, 30, 45, or 60 min. Expression levels were normalized using *ACT2* and *TIP41* as control genes. Data are means ± SEM (n = 3, corresponding to 3 independent pools of 30 seedlings each) for one representative experiment out of two. (b) Kinase activity of MPK7 after immunoprecipitation (IP) with specific antibodies from N-depleted seedlings of the indicated genotypes before (−) and after resupply with 3 mM KNO_3_ or KCl for 15 min.

We then asked whether MPK7 activation in response to nitrate resupply was affected in the *nlp* mutants. Indeed, nitrate-induced MPK7 activation was slightly reduced in the *nlp7-1* and *nlp2-1* single mutants, whereas nitrate-induced MPK7 activation in the *nlp6-1* mutant was similar to that in the WT (Figure 6b). However, the nitrate-induced activation of MPK7 strongly decreased when *nlp7-1* was combined with *nlp2-1* or *nlp6-1*, pointing to functional redundancy among NLP family members. Overall, these results reveal that NLPs play a major role in the activation of the MAPK module.

### The transcription factor NLP7 is not phosphorylated in response to the activation of the MAPK cascade by nitrate

We asked if nitrate-triggered MAPK activation is part of a feedback or feedforward loop by testing whether NLP7 can be a substrate for MPK7 when activated in response to nitrate resupply. We exposed seedlings of Col-0 and the *MPK7-HA* lines to N-free conditions for 3 days prior to resupply with KCl or nitrate for 15 min. A kinase assay using immunoprecipitated MPK7-HA against MBP as a substrate confirmed that MPK7 is activated by nitrate (Figure 7). However, when we used recombinant glutathione S-transferase (GST)-NLP7 purified from bacteria as a substrate, we detected no phosphorylation. As a positive control, we used CPK10-HA and CPK32-HA immunoprecipitated from their respective transgenic lines (Boudsocq et al., 2012); in both cases, the precipitated CPK was able to phosphorylate MBP and GST-NLP7, as previously reported (Liu et al., 2017). Immunoblotting with anti-HA antibodies confirmed that MPK7, CPK10, and CPK32 were present at similar levels across all reactions. These results suggest that MPK7 does not phosphorylate NLP7.

**Figure 7.**
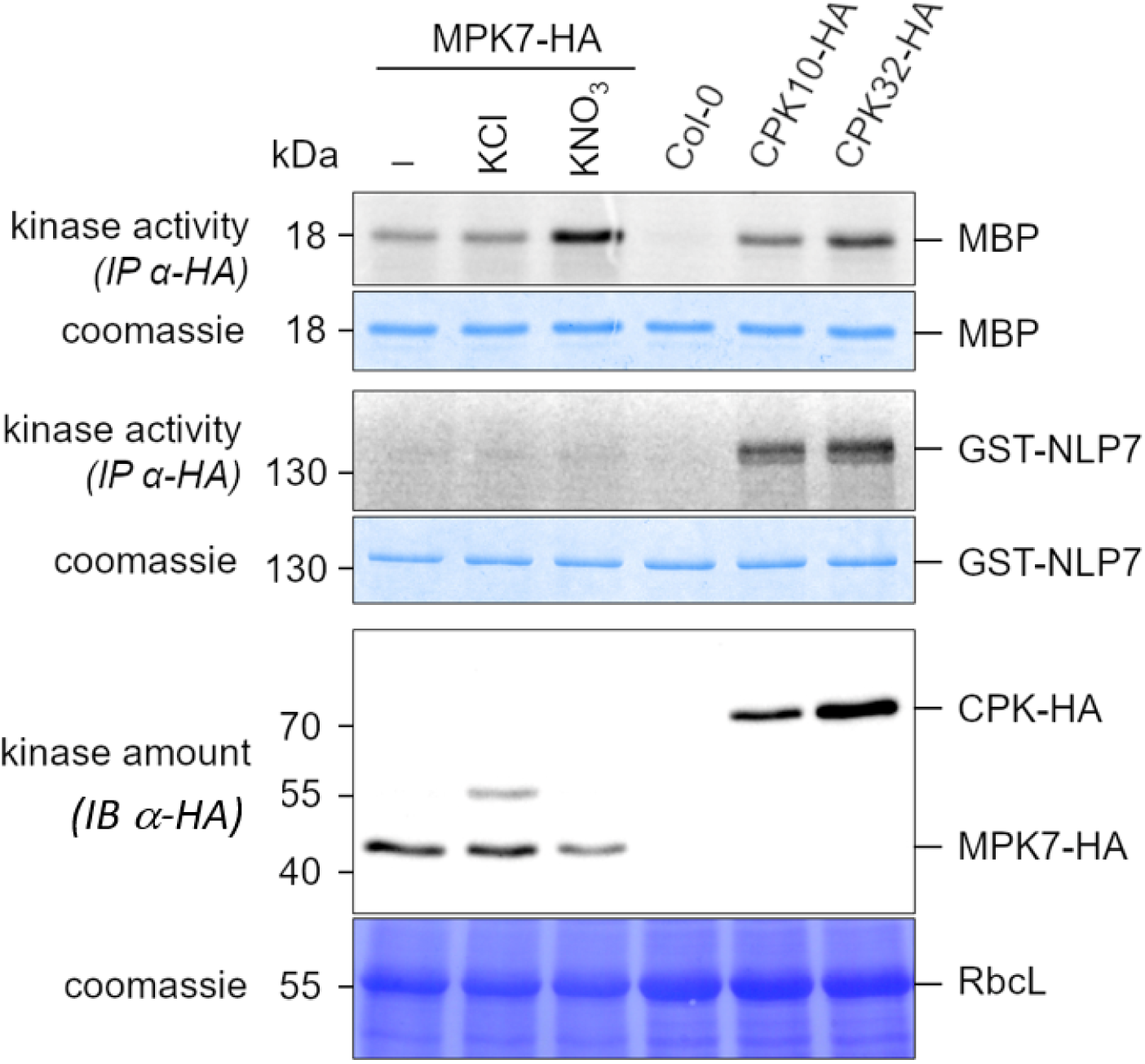
NLP7 is not phosphorylated by the MAP3K13/14–MKK3–MPK7 cascade. Kinase activity of MPK7-HA was monitored using myelin basic protein (MBP) or glutathione S-transferase (GST)-NLP7 as substrates after immunoprecipitation with anti-HA antibody from N-depleted seedlings before (−) or after resupply with 3 mM KCl or KNO_3_ for 15 min (top panels). Anti-HA immunoprecipitates from Col-0 and *35S:CPK-HA* seedlings grown in standard conditions were used as negative and positive controls, respectively. The same protein extracts were immunoblotted with anti-HA antibody to monitor protein abundance (lower panel). Gels and blots were stained with Coomassie to ensure equal loading. RbcL, Rubisco large subunit.

### Nitrate-induced *BT2* expression is impaired by loss of function of the MAP3K13/14– MKK3 module

To unravel the role of the nitrate–NLP2/7–MAP3K13/14–MKK3–MPK7 module in plant responses to nitrate, we asked whether nitrate-regulated gene expression was modified in loss-of-function mutants. We performed RT-qPCR analysis of nitrate-response marker genes in seedlings of Col-0, the two *map3k13 map3k14-cr* double mutants, and two *mkk3* mutants after nitrate resupply for up to 45 min following three days of N depletion, thus using the same experimental conditions as for the MAP kinase activity assays. We quantified the expression of 17 nitrate response marker genes: *NITRATE REDUCTASE 1* (*NIA1*), *NIA2*, *NIR*, *FERREDOXIN-NADP^+^ OXIDOREDUCTASE 2* (*FNR2*), *UROPORPHYRIN METHYLASE 1* (*UPM1*), *NRT2.1*, *CHLORIDE CHANNEL-A* (*CLCA*), *GLUCOSE-6-PHOSPHATE DEHYDROGENASE* (*G6PDH*), *BTB AND TAZ DOMAIN PROTEIN 2* (*BT2*), *HRS1*, *HRS1 HOMOLOG 1* (*HHO1*), *CIPK3*, *TGACG MOTIF-BINDING FACTOR 1* (*TGA1*), *TGA4*, *LOB DOMAIN-CONTAINING PROTEIN 37* (*LBD37*), *LBD38*, and *LBD39*. Among this subset of nitrate-regulated genes, *BT2* expression was induced to a level approximately ∼30% less at 30 and 45 min after nitrate resupply in the two *map3k13 map3k14-cr* double mutants, and approximately ∼50% less at 45 min in both *mkk3* mutants, compared to the respective WT plants (Figure 8a, b). None of the other genes showed differential expression in *map3k13 map3k14-cr* or *mkk3* mutants in response to nitrate resupply (Figure S5). Hence, the MAP3K13/14–MKK3–MPK7 module is involved in the nitrate-dependent induction of *BT2* expression.

**Figure 8.**
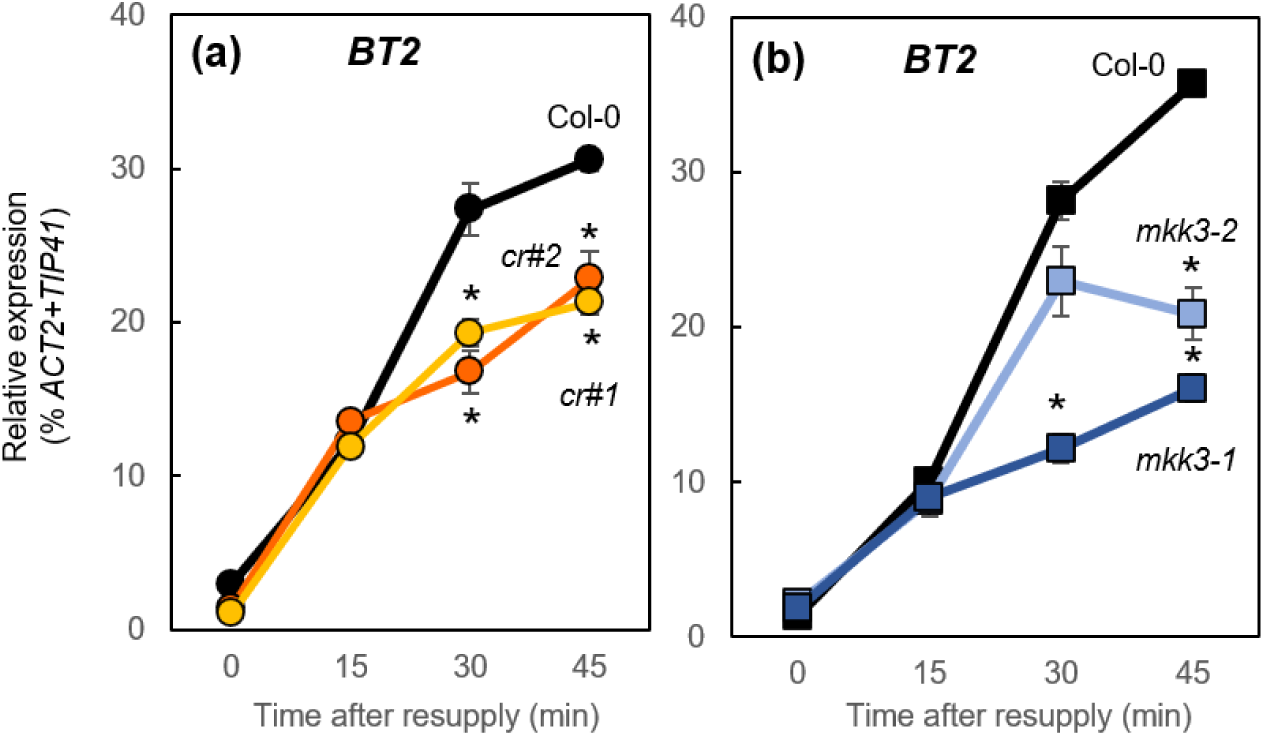
Nitrate-induced *BT2* expression is regulated by the MAP3K13/14–MKK3 module. Relative *BT2* transcript levels after nitrate addition to N-starved seedlings. Seedlings were cultivated and treated as in Figure 1a. Expression was measured by RT-qPCR and normalized using *TIP41* (At4g34270) and *ACTIN2* (*ACT2*, At3g18780) as control genes. Data are means ± SEM (n = 3, corresponding to 3 independent batches of 30 seedlings each) of one representative experiment out of three. Asterisks indicate statistically significant differences between the wild-type Col-0 and mutants (*P* < 0.05, Student’s *t*-test). (a) Relative *BT2* transcript levels in Col-0, *map3k13 map3k14*-cr#*1 (cr#1)*, and *map3k13 map3k14*-cr#*2 (cr#2)*. (b) Relative *BT2* transcript levels in Col-0, *mkk3-1*, and *mkk3-2*.

### Nitrate uptake is enhanced by the loss of MAP3K13 and MAP3K14 function

BT2 regulates nitrate uptake (Araus et al., 2016). To test if the loss of the nitrate–NLP2/7– MAP3K13/14–MKK3–MAPK module affects nitrate uptake, we measured high-affinity and low-affinity ^15^N nitrate influx in WT and mutant seedlings grown under constant nitrate-replete conditions (3 mM) for 11 days. High-affinity nitrate influx increased by 13 and 22%, respectlively, in the *map3k13 map3k14-cr* mutant lines compared to WT. However, we observed no significant difference in the *mkk3* mutants relative to WT (Figure 9a). Low-affinity nitrate influx was similar among all genotypes (Figure 9b).

**Figure 9.**
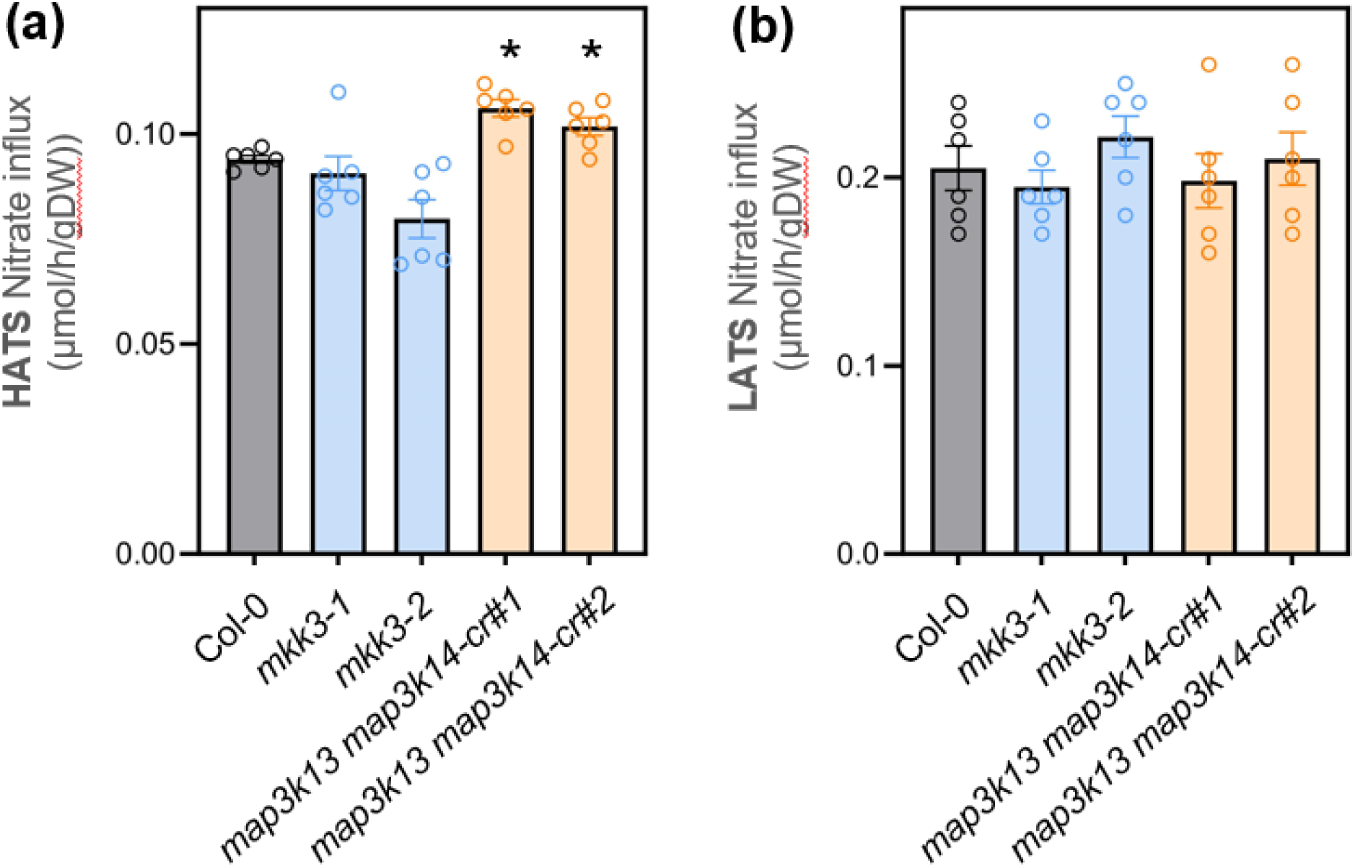
The *map3k13 map3k14* mutants show increased high-affinity nitrate uptake. Seedlings were grown vertically on medium containing 3 mM nitrate for 11 days. ^15^NO ^−^ influx was measured after 5 min of labeling in complete nutrient solution containing 0.2 mM or 6 mM ^15^NO ^−^. (a) High-affinity transport system (HATS), measured using 0.2 mM nitrate. (b) Low-affinity transport system (LATS), measured using 6 mM nitrate (and subtraction of HATS). Data are means ± SEM from six replicates corresponding to five seedlings each. One of two independent experiments is shown. Asterisks indicate statistically significant differences between the WT (Col-0) and mutants (*P* < 0.01, Student’s *t*-test). DW, dry weight.

We then asked whether this subtle increase in high-affinity nitrate influx in *mapk3K13 map3k14-cr* seedlings grown under nitrate-replete conditions would lead to changes in plant growth and nitrate content. Accordingly, we measured rosette and root biomass, as well as nitrate contents, of plants grown under the same conditions as for the nitrate influx measurements or under a low nitrate supply (0.2 mM). We detected no significant difference in biomass or nitrate content under any of these conditions or among genotypes (Figure S6). Thus, the loss of MAP3K13 and MAP3K14 function affected *BT2* expression and high-affinity nitrate uptake, but without modifying biomass or nitrate content under our experimental conditions.

### Overexpression of *MAP3K13* modifies low N-triggered senescence in a MKK3-dependent manner

Since we did not identify any clear nitrate-related phenotype in loss-of-function mutants for components of the MAPK cascade, we turned to a gain-of-function approach and generated lines overexpressing *MAP3K13* under the control of the strong *UBIQUITIN10* promoter (*MAP3K13*-OE #1, #2). *MAP3K13* expression levels in these lines increased up to 38-fold compared to the WT (Figure S7a). We first asked whether MPK7 activity and its stimulation by nitrate were modified in these OE lines under our standard nitrate resupply conditions. We observed a slight increase in MPK7 activity between WT and OE lines (Figure S7b). To unravel the consequences of *MAP3K13* overexpression on plant responses to N availability, we imposed severe N starvation by growing plants under a short-day photoperiod under limiting N supply (0.5 mM nitrate) for 38 days, followed by 32 days under a long-day photoperiod. At this developmental stage, the plants had flowered and their growth was strongly limited by the low N supply, as also evidenced by the loss of chlorophyll and the accumulation of anthocyanins (Figure 10a). Under these conditions, we observed a difference in rosette color in two independent *MAP3K13*-overexpressing lines relative to WT and to *mkk3-1* (Figure 10a) due to a strong decrease in chlorophyll levels (Figure 10b) and increased anthocyanin accumulation (Figure 10c).

**Figure 10.**
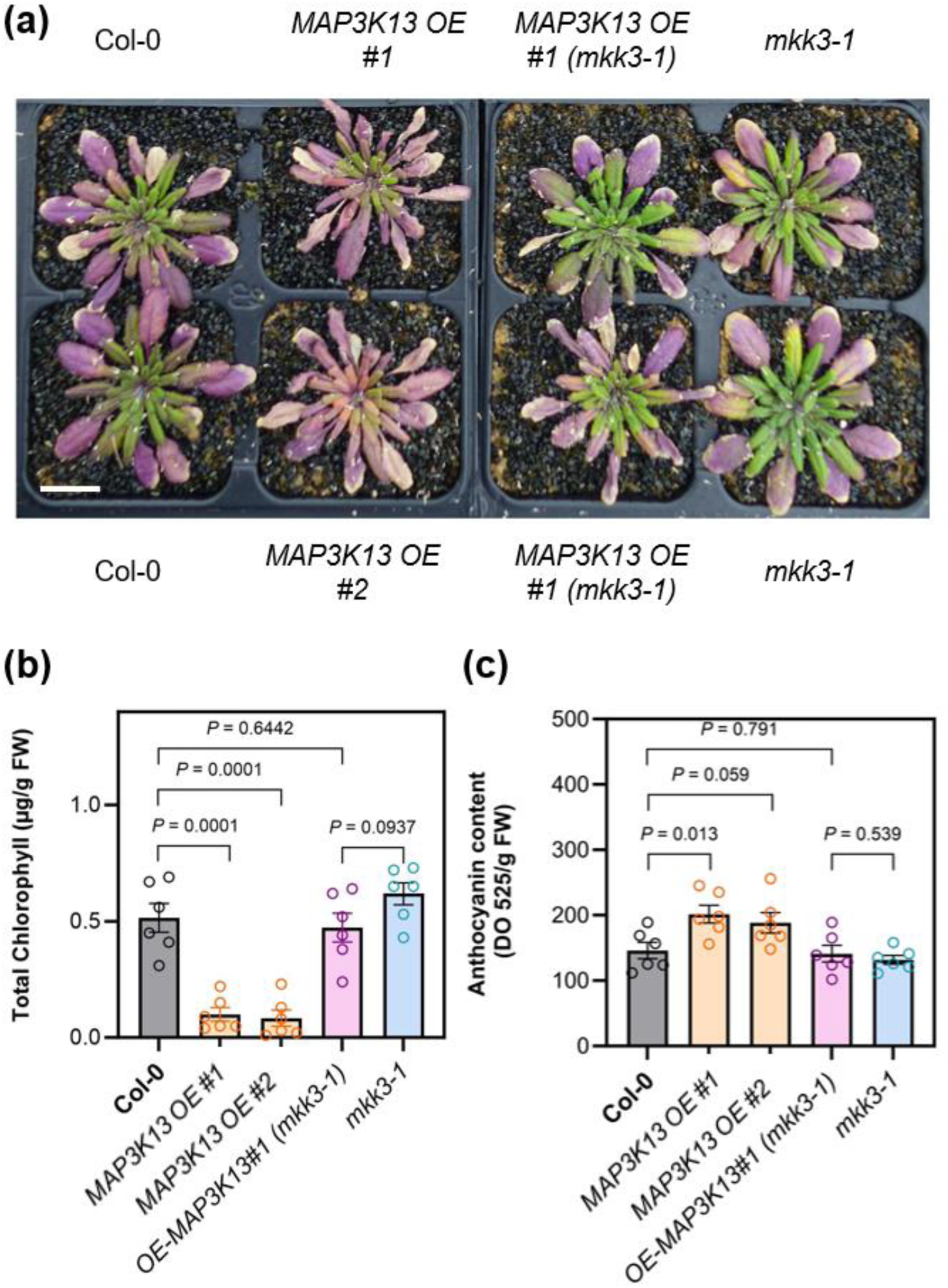
Chlorophyll depletion and anthocyanin accumulation in *MAP3K13* overexpressor lines after severe N limitation is dependent on MKK3. (a) Top view of 70-d-old rosettes from plants grown under limiting (0.5 mM) nitrate supply. Flower stalks were cut off prior to taking the photographs. (b) Total chlorophyll contents in rosettes. (c) Anthocyanin contents in rosettes. Data are means ± SEM (n = 5, each sample corresponding to one plant and shown as a circle). *P*-values were determined by Student’s *t*-test. Scale bar, 2 cm.

To confirm that the overexpression of *MAP3K13* affected chlorophyll and anthocyanin levels via the MAP3K13–MKK3 module, we introgressed *mkk3-1* into both *MAP3K13*-OE lines. Importantly, the loss of MKK3 function abolished the changes in chlorophyll and anthocyanin contents caused by *MAP3K13* overexpression (Figure 10a, b, c). Taken together, these observations confirm (by genetic means) that MKK3 functions downstream of MAP3K13 and suggest that this module may be involved in regulating N depletion–stimulated senescence.

## DISCUSSION

Kinase- or phosphatase-mediated protein phosphorylation or dephosphorylation enables the rapid and specific initiation of signal transduction, as well as the amplification of signals in response to external stimuli (Jonak et al., 2002; Zhang and Zhang, 2022). Among the myriad of kinases found in living cells, MAPK cascades have been implicated in responses to environmental cues in all eukaryotic organisms (Cargenello and Roux, 2011; Xie et al., 2023). Since their discovery in plants in 1993, MAPK cascades have been shown to function mainly in stress perception and development (Xu et al., 2015; Zhang et al., 2018). However, the importance of MAPKs for nutrient signaling has been unclear (Chardin et al., 2016). In this study, we identified a MAP3K13/14–MKK3–MPK7-dependent signaling pathway that is induced within minutes by nitrate in an NLP-dependent manner and is involved in plant responses to N availability.

### A nitrate-activated MAPK pathway

Few previous studies have linked MAPKs to N signaling pathways. Forde et al. (2013) showed that MEKK1 is involved in regulating root architecture in response to external glutamate, although the upstream and downstream kinases and their target proteins have yet to be discovered. Here, we showed that nitrate resupply to N-starved seedlings and nitrate addition to ammonium-grown seedlings triggered the activation of the group C MAPK MPK7. Indeed, the nitrate-triggered activation of MPK7 occurred very rapidly and was highly sensitive to nitrate and independent of nitrate reduction by nitrate reductase. As nitrate reductase not only catalyzes the first step of nitrate assimilation into organic N compounds, but is also a major producer of NO, this finding suggests that the induction of the MAP3K13/14–MKK3–MPK7 module is nitrate-specific and independent of NO signaling. However, this hypothesis needs to be empirically tested using NO donors and scavengers. Interestingly, our results also suggest that the nitrate sensor function of the nitrate transceptor NPF6.3 is not required for the nitrate-triggered activation of MPK7. Thus, nitrate sensing by NLPs, as discovered recently (Liu et al., 2022), might be the first event triggering this MAPK cascade.

### *MAP3K13* and *MAP3K14* transcripts rapidly accumulate in response to nitrate resupply

MAP3K13 and MAP3K14 belong to the subclade III of MEKK-like MAP3Ks, whose functions have only recently been partially unraveled (Colcombet et al., 2016). In this study, we were interested in *MAP3K13* and *MAP3K14* due to their strong NLP7-dependent upregulation in N-depleted seedlings in response to nitrate resupply (Marchive et al., 2013). In fact, nitrate was shown to regulate *MAP3K13* and *MAP3K14* expression in many studies (Wang et al., 2003; Scheible et al., 2004), while the induction of MAP3K19 expression appeared to depend on the experimental conditions (Liu et al., 2020). Additionally, like the MAP*3K13* and *MAP3K14* promoters, the *MAP3K19* promoter was one of the 851 genomic regions bound by NLP7 (Marchive et al., 2013). However, we detected no differential expression of *MAP3K19* in response to nitrate in this earlier study, suggesting the involvement of additional players regulating *MAP3K19* expression. Importantly, we showed that the loss of MAP3K13 and MAP3K14 function completely abolished nitrate-triggered MPK7 activation, indicating that MPK7 activation strictly depends on these two MAP3Ks under our experimental conditions. Nevertheless, we cannot exclude the possibility that other nitrate-regulated subclade III MAP3Ks might be involved in regulating physiological traits in Arabidopsis in response to N availability and that they act redundantly with MAP3K13 and MAP3K14 despite a much lower accumulation of their encoding transcripts in response to nitrate resupply. Interestingly, it was recently reported that, in parallel to MAP3K13 and MAP3K14, MAP3K19 and MAP3K20 play key roles in nitrate and light activation of the module in seeds to break secondary dormancy (Regnard et al., 2024).

### NLP transcription factors are upstream regulators of the MAP3K13/14–MKK3–MPK7 module

We further determined that the very rapid, highly transient induction of *MAP3K13* and *MAP3K14* expression in response to nitrate resupply was compromised in mutants lacking function of the key nitrate response transcription factors NLP7 and NLP2. NLP7 and NLP2 govern a large part of the gene regulatory network in response to nitrate (Marchive et al., 2013; Konishi et al., 2021; Alvarez et al., 2020; Durand et al., 2023). Using single and double mutants, we showed that MPK7 activation was strongly reduced in *nlp2-1 nlp7-1* and *nlp6-1 nlp7-1* double mutants, whereas the loss of either NLP7, NLP6, or NLP2 at best only slightly affected MPK7 activation. These results resemble those shown for nitrate-regulated gene expression in the same mutant backgrounds (Castaings et al., 2009; Marchive et al., 2013; Konishi et al., 2021; Alvarez et al., 2020; Durand et al., 2023; Cheng et al., 2023), suggesting partial redundancy among NLPs. The analysis of higher-order mutants of *NLP* genes should help address this issue. In the context of the nitrate-induced activation of the MKK3 module in seeds, in addition to NLP8, which governs nitrate-dependent primary seed dormancy (Yan et al., 2016), NLP7 and NLP9 play redundant roles in secondary seed dormancy (Regnard et al., 2024). In addition to the transcriptional regulation of *MAP3K13* and *MAP3K14* by NLPs, other regulatory mechanisms might also be involved. For example, MAP3K18 undergoes post-translational regulation by protein degradation (Mitula et al., 2015; Zhou et al., 2021). Interestingly, MAP3K13 and MAP3K14 were highly unstable when their encoding constructs were expressed in protoplasts (Sözen et al., 2020), suggesting that a similar type of protein degradation may occur.

### MAP3K13 and MAP3K14 function in nitrate-regulated gene expression and in the regulation of nitrate uptake and N limitation–triggered senescence

We showed that the loss of function of MAP3K13 and MAP3K14 or MKK3 impaired, but did not abolish, the nitrate-triggered accumulation of transcripts of *BT2*, a Bric-a-Brac/Tramtrack/Broad (BTB) family gene, when nitrate was resupplied to N-depleted seedlings. However, the expression of the other classical PNR marker genes tested here did not change in the mutants in response to nitrate. Indeed, *BT2* was previously identified as the most central and most highly connected gene in the N use efficiency (NUE) network (Araus et al., 2016) and is a direct target of NLP7 (Sato et al., 2017; Durand et al., 2023). A transcriptome deep sequencing approach could help unravel whether the nitrate-mediated regulation of other genes is controlled by the MAP3K13/14–MKK3–MPK7 cascade. However, another study showed that the MAP3K18–MKK3–MPK7 cascade has a modest effect on gene expression (Danquah et al., 2015). BTB proteins drive selective protein ubiquitination via their assembly into Cullin3-based ubiquitin ligases (Figueroa et al., 2005; Mandadi et al., 2009; Zhao et al., 2016; An et al., 2018). Using a reverse genetics approach, Araus and coworkers showed that nitrate uptake under low-nitrate conditions increased in the *bt1 bt2* double mutant compared to WT (Araus et al., 2016). Here, high-affinity nitrate uptake slightly increased in the *map3k13 map3k14* double mutants when plants were grown under constant nitrate-replete conditions. This result is consistent with the notion that BT2 acts as a repressor of nitrate transport; however, our results differ from those obtained by Araus et al. (2016) showing increased nitrate uptake in the *bt1 bt2* double mutant under limiting N supply. Clearly, the regulatory mechanisms involving the MAPK cascade in the response to N availability are still far from understood and might exceed the consequences of the downregulation of *BT2* expression.

### *MAP3K13* overexpression accelerates N limitation-triggered senescence

BT2 was recently shown to repress anthocyanin biosynthesis in apple (*Malus domestica*) (An et al., 2020). We found that under severe N limitation-triggered senescence, anthocyanin accumulation and chlorophyll loss were amplified in Arabidopsis lines overexpressing *MAP3K13*, a trait that was dependent on MKK3. In apple, the effect of BT2 on anthocyanin accumulation is mediated by a BT2–Teosinte branched1/CYCLOIDEA/PROLIFERATING CELL FACTOR (TCP)46–MYB1 protein module that regulates the degradation of MYB1. Interestingly, a TCP-type transcription factor (TCP20) participates in the gene regulatory network in response to nitrate via its interaction with NLP6 and NLP7 (Guan et al., 2017), and thus a similar module might contribute to the stability of NLP transcription factors. However, other regulatory proteins contribute to the regulation of anthocyanin biosynthesis in response to N starvation, such as LBD transcription factors (Rubin et al., 2009) and the E3 ubiquitin ligase NITROGEN LIMITATION ADAPTATION (NLA) (Peng et al., 2007; 2008). MKK3-modulated senescence was also revealed in other studies. Indeed, *MAP3K18* overexpressors showed accelerated leaf senescence (Matsuoka et al., 2015). As leaf senescence is controlled by ABA, among other regulators (Song et al., 2016), this observation is in agreement with the notion that the MAP3K18–MKK3–MPK7 module functions in ABA signaling (Danquah et al., 2016). More recently, group C MAPKs were shown to regulate ABA-induced leaf senescence (Li et al., 2024), and ABA was suggested to trigger the MAP3K17/18–MKK3–group C MAPK cascade, resulting in the phosphorylation of MBD10, a member of the methyl-CpG-binding domain protein family (Grafi et al., 2007). In addition, the MKK4/5–MPK1/2 cascade is involved in salicylic acid (SA)-regulated senescence (Zhang et al., 2020), suggesting that group C MAPKs contribute to both ABA- and SA-regulated senescence. Here we found that the altered senescence in *MAP3K13* overexpressors in triggered by N limitation was dependent on MKK3. This finding indicates that the SA–MKK4/5 module is not involved in the MAP3K13/14-regulated senescence mechanism. The mechanism underlying the regulation of senescence by the MAP3K13/14–MKK3–MPK7 module requires further studies.

### Do some roles of MAP3K13 and MAP3K14 bypass MKK3–MPK7?

Intriguingly, whereas the characterization of nitrate-induced MPK7 activation in mutants lacking function of MAP3K13 and MAP3K14 or MKK3 suggested that they function in the same cascade, their phenotypes only partially overlapped. Thus, MAP3K13 and MAP3K14 may have MKK3-independent functions. Indeed, in addition to their well-known function in catalyzing phosphotransfer to downstream kinases in signaling cascades, other molecular functions have been described for MAP3Ks. For example, the subclade II MAP3K MEKK1 interacts with the transcription factor WRKY53 and binds to the *WRKY53* promoter to regulate its expression (Miao et al., 2007). Similarly, the RAF-type MAP3K-like VH1-INTERACTING KINASE (VIK) bypasses a standard MAPK cascade and phosphorylates and regulates a sugar transport protein (Wingenter et al., 2012). However, RAF kinases, which were occasionally shown to function like MAP3Ks in animal systems, do not clearly function in this manner in plants (Wang, 2024). MAPK3K13 and MAP3K14 may also be involved in a second MAPK cascade through a distinct MAP2K. For example, MAP3K20, which also belongs to subclade III of the MAP3K family together with MAP3K13 and MAP3K14, may act upstream of MKK5–MPK6 and MKK3 (Li et al., 2017; Sözen et al., 2020; Bai et al., 2022). To test these hypotheses, it would be interesting to characterize the MKK3-independent roles and molecular mechanisms by further comparing the phenotypes of plants impaired in MKK3 function and plants impaired in MAP3K13 and MAP3K14, as well as to identify the targets and interactors of MAP3K13 and MAP3K14, for example by performing phosphoproteomics and identifying interacting proteins.

### What are the target proteins of the module?

To further understand the biological role of this nitrate-triggered MAPK cascade in seedlings and adult plants facing fluctuations in nitrate availability, it will be crucial to identify the target proteins of these protein kinases. Transcription factors are often the downstream effectors of signaling cascades including MAPK cascades. Here we showed that members of the NLP family of transcription factors are upstream regulators of the nitrate-mediated increase in *MAP3K* transcription and in the activation of the cascade. Feedback regulation via phosphorylation of NLP7 by MPK7 would be an attractive mechanism to fine-tune nitrate signaling. However, we did not find evidence for MPK7-dependent phosphorylation of NLP7. It was previously suggested that nitrate reductase could be phosphorylated by MAPKs (Popescu et al., 2008; Wang et al., 2010; Hoehenwarter et al., 2013). The link between the nitrate signaling pathway and MBD10, a recently identified substrate of group C MAPKs (Li et al., 2024), needs to be investigated further, as other MBDs repress gene expression by recognizing methylated DNA (Ichino et al., 2022). Furthermore, the transcription factor ERF4 was identified as a target of group C MAPKs involved in the release of seed dormancy in Arabidopsis (Chen et al., 2023). Chen and colleagues showed that MPK7 phosphorylates ERF4, leading to its degradation and thus releasing the repression of *alpha-EXPANSIN* (*EXP*) expression. Expansin-induced cell wall extension modulates plant growth (Cosgrove, 2024), and thus a similar module may be involved in growth stimulation by nitrate. Indeed, *EXP7* expression is regulated by nitrate (Canales et al., 2014).

In addition to identifying of the targets of this MAPK cascade during early nitrate signaling, studying the crosstalk with other signaling pathways involving the MKK3–group C MAPKs module will help dissect the molecular mechanisms by which this previously unknown MAPK signaling pathway contributes to the highly controlled, rapid responses of plants to nitrate exposure. The study of nitrate signaling pathways is highly relevant to understanding the regulation of nitrate assimilation and transport in order to improve NUE in crops under sustainable farming conditions.

## EXPERIMENTAL PROCEDURES

### Plant materials

All Arabidopsis (*Arabidopsis thaliana*) materials were in the Columbia (Col-0) background. The mutants *mkk3-1* (SALK_051970), *mkk3-2* (SALK_208528), *nlp7-1* (SALK_26134), *nlp6-1* (SALK_018362), and *nlp2-1* (SAIL_139_D05) were previously described (Dóczi et al., 2007; Sözen, et al., 2020; Castaings et al., 2009; Marchive et al., 2013; Konishi et al., 2021). All mutant seeds were obtained from the Eurasian Arabidopsis Stock Centre (uNASC), and the plants were backcrossed twice to Col-0. The double mutant*s nlp2-1 nlp7-1 and nlp6-1 nlp7-1* were described in Durand et al. (2023) and Cheng et al. (2023), respectively. The *chl1-5* and *chl1-9* mutant lines were obtained from Dr. Y.F. Tsay (Tsay et al., 1993; Ho et al., 2009) and *the nia1 nia2* mutant line (also referred to as G’43, Wilkinson and Crawford, 1993) from the Eurasian Arabidopsis Stock Centre (uNASC). Transgenic lines expressing HA-tagged MPK7 using a construct containing the entire *MPK7* locus including its promoter were described in Sözen et al. (2020); the *mkk3-1* mutant line complemented by *MKK3-YFP* using a construct containing the entire *MKK3* locus including its promoter was described in Danquah et al. (2015); the *35S:CPK10-HA* and *35S:CPK32-HA* lines were described in Boudsocq et al. (2012).

The *map3k13 map3k14* double mutants were generated in parallel to the genome-edited *map3k14–cr* mutants used in Sözen et al. (2020). Plants were transformed with the plasmid pDGE65-MAP3K14-CR by the floral dip method (Clough and Bent, 1998); positive primary transformants were genotyped by sequencing for mutations at both the *MAP3K13* and *MAP3K14* loci, as the CRISPR/Cas sgRNA targeted both loci. The pDGE65-MAP3K14-CR transgene was removed by segregation; two lines derived from independent transformation events and carrying homozygous mutations in *MAP3K13* and *MAP3K14* were selected. These lines are referred to as *map3k13 map3k14-cr#1* and *map3k13 map3k14-cr#2*. To obtain the corresponding single mutants, the *map3k13 map2k14-cr#1* double mutant was crossed to Col-0, and the F2 progeny were genotyped to identify WT plants and single and double mutants.

To produce *MAP3K13* overexpressing lines (OE), the full-length *MAP3K13* coding sequence was cloned into the pDNR207 vector as described previously (Sözen et al., 2020) using the primers listed in S1. The Sanger sequencing-verified pDNR207 vector carrying the *MAP3K13* coding sequence with a stop codon was recombined in pUB-dest (Grefen et al., 2010) using Gateway technology following the manufacturer’s instructions to generate pUB-MAP3K13. This vector was transformed into Agrobacterium (*Agrobacterium tumefaciens*) strain C58C1 and used to transform Col-0 by the floral dip method. Primary transformants were selected on Estelle and Sommerville medium (Estelle and Sommerville, 1987) containing 20 μg L^−1^ of hygromycin B. In the subsequent generations after selfing, homozygous lines were selected following prior selection for a single T-DNA insertion event. In addition, the transgene was introduced into the *mkk3-1* mutant by genetic crossing, and seeds homozygous for *mkk3-1* and the transgene were used for analysis.

### Growth conditions

For nitrate response assays to measure mRNA levels and kinase activity, seeds were surface sterilized with 15% (v/v) commercial bleach in ethanol for 10 min, washed 5 times with sterile water, stratified for 3 days at 4°C in the dark, and grown in liquid medium containing 0.07% (w/v) MES pH 5.8, 1 mM KH_2_PO_4_, 0.5 mM MgCl_2_, 0.5 mM CaSO_4_, microelements as described in Estelle and Somerville (1987), 0.25% (w/v) sucrose, and 3 mM KNO_3_ for 11 days under a long-day photoperiod (16-h light/8-h dark) at 25°C at a light intensity of 80 µmol·m^−2^·s^−1^ provided by Philips F17T8/TL741 light sources. After 7 days of growth, the medium was replaced every 2 days with fresh medium. Eleven-day-old seedlings were subjected to N-limited conditions for 3 days using fresh medium containing 3 mM KCl instead of 3 mM KNO_3_. After 3 days of N starvation, 3 mM nitrate was added for the indicated duration, and the seedlings were harvested at several time points after nitrate addition (at 3–5 h after dawn).

To analyze adult plants, seeds were stratified for 3 days at 4°C and grown in sand culture supplemented with nutrient solution (Loudet et al., 2003) containing either 0.5 mM nitrate (limiting nitrate supply) or 5 mM nitrate (non-limiting nitrate supply), both supplied as a mixture of KNO_3_ and Ca(NO_3_)_2_. For N limitation conditions, ionic equilibrium of the medium was ensured by partly replacing KNO_3_ and Ca(NO_3_)_2_ by KCl and CaSO_4_. Plants were kept under a short-day photoperiod (8-h light/16-h dark) for 38 days before being transferred to a long-day photoperiod (16-h light/8-h dark) for 32 days of growth. The growth conditions were 21°C/18°C during day and night, respectively, with 65% relative humidity and 150 μmol·m^−2^s^−1^ irradiation provided by Osram 36w light sources. Rosettes were harvested at 3–4 h after dawn for all analyses (anthocyanin, chlorophyll, and nitrate contents). The tissues were flash-frozen in liquid nitrogen and stored at −80°C until use.

### Gene expression analysis

Measurements of relative transcript levels were performed on seedlings (1.02 developmental growth stage according to Boyes et al. (2001)) cultivated in liquid medium as described above. Each sample comprised 26–34 seedlings that were flash-frozen after harvest as described above. Total RNA was extracted from the samples using an RNeasy Plant Mini Kit (QIAGEN). First-strand cDNAs were synthesized according to Daniel-Vedele and Caboche (1993) using Moloney murine leukemia virus reverse transcriptase and oligo(dT)15 primers (Promega) or using a SuperScript III Fast strand synthesis system (Invitrogen). cDNAs were first checked by PCR for contamination with genomic DNA using primers amplifying *EF1a* (At5g60390) flanking introns, as listed in Table S1, and relative expression was measured in 384-well plates with a QuantStudio 5 System device (40 3-step cycles; step 1, 95°C, 5 s; step 2, 58°C, 30 s; step 3, 72°C, 30 s) using a Takyon qPCR kit for SYBR assay (Eurogentec) (2.5 µL TAKYON SYBER 2X, 0.015 µL forward primer [100 µM], 0.015 reverse primer [100 µM], 2.5 µL cDNA diluted 1:20 [v/v] per well). Relative expression was calculated according to the 2^−ΔCt^ method (Livak et al., 2001). Target gene expression was normalized to the geometric mean of the expression of two plant genes, either *APT* (At1g27450) and *ACTIN2* (*ACT2*, At3g18780) or *TIP41 (*At4g34270) and *ACTIN2.* All primer sequences are listed in Table S1.

### Kinase assays and immunoblotting

Kinase assays using anti-MPK7 (Doczi et al., 2007), anti-MPK2 (Ortiz-Masia et al., 2007), anti-HA (Sigma-Aldrich H6908), and anti-MPK1 (Ortiz-Masia et al., 2007) antibodies were performed as described by Danquah et al. (2015) and Ortiz-Masia et al. (2007). It should be noted that the anti-MPK2 antibodies can specifically immunoprecipitate MPK2, although they fail to detect the protein on immunoblots of total proteins (Ortiz-Masia et al., 2007). A detailed protocol can be found in the supplemental data of Sözen et al. (2020). In some cases, protein levels were monitored by immunoblot analysis following Bio-Rad recommendations. Briefly, proteins were separated on 10% (w/v) SDS-PAGE gels and transferred onto polyvinylidene difluoride membranes (Bio-Rad). The membranes were blocked in 5% (w/v) nonfat dry milk and probed with anti-HA antibodies (Roche 11867431001; 1:10,000 dilution), followed by horseradish peroxidase-coupled secondary anti-rat antibody (Sigma-Aldrich A9037, 1:10,000 dilution). Horseradish peroxidase activity was detected with a Clarity Western ECL Substrate Reaction kit (Bio-Rad) and a ChemiDoc Imaging System (Bio-Rad). The blots were stained with Coomassie Brilliant Blue for protein visualization.

### NLP7 phosphorylation assay

The full-length coding sequence of *NLP7* was cloned into the vector pGEX-2T (GE Healthcare) as a BamHI-StuI fragment. Recombinant GST-NLP7 was produced in *Escherichia coli* BL21(DE3)pLys cells (Stratagene) and purified as previously described (Boudsocq et al, 2012). For the kinase assay, seedlings were ground in liquid nitrogen and homogenized in immunoprecipitation buffer (50 mM Tris-HCl pH 7.5, 5 mM EDTA, 5 mM EGTA, 50 mM β-glycerophosphate, 10 mM sodium fluoride, 1 mM orthovanadate, 2 mM DTT, 1X anti-protease cocktail [Roche], 150 mM NaCl, 1% [v/v] Triton X-100) prior to centrifugation at 21,100 g for 15 min at 4°C. Protein concentration was determined by the Bradford method. Protein extract (200 μg) was incubated with 1 μL polyclonal anti-HA antibody (Sigma-Aldrich H6908) in immunoprecipitation buffer for 2 h at 4°C. After adding 20 μL of a 50% [w/v] slurry of Protein A-Sepharose beads, the sample was incubated for another 1 h at 4°C. The immunoprecipitates were washed three times in immunoprecipitation buffer and twice in protein kinase buffer (20 mM Tris-HCl pH 7.5, 10 mM MgCl_2_, 1 mM DTT, 1 mM CaCl_2_). The immunoprecipitates were incubated in reaction buffer (20 mM Tris-HCl pH 7.5, 1 mM CaCl_2_, 10 mM MgCl_2_, 1 mM DTT, 50 μM cold ATP, 2 μCi [γ-33P] ATP) with 1 µg substrate (purified GST-NLP7 or myelin basic protein (MBP, Sigma M1891)) at room temperature for 30 min. The reaction was stopped by adding SDS-PAGE loading buffer. The samples were heated at 95°C for 3 min and separated by SDS-PAGE. Phosphorylation was detected from the dried gels with a Typhoon imaging system (GE Healthcare). Gels were stained with Coomassie as a loading control.

### Root ^15^N influx measurements

The influx of ^15^NO_3_^−^ into roots was assayed as previously described (Orsel et al., 2004). Seedlings were grown on 3 mM nitrate medium (described above) solidified with 0.8% (w/v) agarose (Sigma A1296) and placed vertically. The seedlings were transferred first to 0.1 mM CaSO_4_ for 1 min and then to complete nutrient solution containing ^15^NO_3_ ^−^ (atom% ^15^N: 99%) at the indicated concentrations for 5 min and finally to 0.1 mM CaSO_4_ for 1 min. The roots were dried for 72 h at 80°C and analyzed using a FLASH 2000 Organic Elemental Analyzer coupled to an isotope ratio mass spectrometer (ThermoFisher Scientific). The influx of ^15^NO ^−^ was calculated based on the total N and ^15^N contents of the roots.

### Nitrate, chlorophyll, and anthocyanin measurements

After ethanolic extraction of flash-frozen and finely ground plant materials, nitrate and chlorophyll contents were determined as in Orsel et al. (2004) and Krapp et al. (1993), respectively. For anthocyanin measurements, plant materials were extracted and analyzed as described in Diaz et al. (2006).

### Statistical analysis

Experiments were performed at least twice using replicates as indicated in the legends. Statistical analyses, as indicated in the legends, were performed with GraphPad Prism 10.0.1 (GraphPad Software, La Jolla, CA, USA) applying a significance threshold of 0.05.

## ACCESSION NUMBERS

Sequence data from this article can be found in the Arabidopsis Genome Initiative or GenBank/EMBL databases under the following accession numbers: At1g10210 (*MPK1*), At1g59580 (*MPK2)*, At2g18170 (*MPK7*), At5g40440 (*MKK3*), At1g07150 (*MAP3K13*), At2g30040 (*MAP3K14*), At2g32510 (*MAP3K17*), At1g05100 (*MAP3K18*), At5g67080 (*MAP3K19*), At3g50310 (*MAP3K20*), and At3g48360 (*BT2*).

## AUTHOR CONTRIBUTIONS

JC and AK designed the experiments, STS, CC, VB, MB, JC, and AK performed the experiments and analyzed the data, AM measured the ^15^N enrichment, JC and AK wrote the first draft, and all authors edited the final manuscript.

## ACKNOWLEDGMENTS

Work in our laboratories was partly funded by MAPKSEED ANR grant ANR14-CE19-007 to MB, JC, and AK. This work has benefited from a French State grant (LabEx Saclay Plant Sciences-SPS ANR-10-LABX-0040-SPS), managed by the French National Research Agency under an “Investments for the Future’’ program (ANR-11-IDEX-0003-02), to CC, AK, SB, and JC. STS was supported by a grant from the AgreenSkills+ EU fellowship program (FP7-609398). We are grateful for stimulating discussions with Dr. Heribert Hirt (KAUST, Saudi Arabia), and we thank Dr. Zsolt Kelemen (IJPB, France) for frequent sample transport between our institutes. This work has benefited from the support of the IJPB Plant Observatory (PO) platforms PO-Plants and PO-Biochem.

## SUPPORTING INFORMATION

Additional Supporting Information may be found in the online version of this article

**Figure S1.** Transcript levels of *MAP3K17*, *MAP3K18*, *MAP3K19*, and *MAP3K20* in response to nitrate resupply.

**Figure S2.** MPK7-HA activation and MPK-HA protein abundance in response to nitrate resupply.

**Figure S3.** MAP3K13 and MAP3K14 are required for MPK7 activation by nitrate.

**Figure S4.** Transcript levels of group C *MPK*s and *MKK3* after nitrate resupply.

**Figure S5.** The expression of many nitrate-regulated marker genes is not modified in *map3k13 map3k14-cr* or *mkk3* mutants compared to the wild type.

**Figure S6**. Biomass and nitrate contents of mutants grown under nitrate-replete conditions or limiting nitrate supply are not affected by the loss of MAP3K13 and MAP3K14 or MKK3 function.

**Figure S7**. Molecular characterization of *MAP3K13* overexpressing lines.

**Table S1.** Nucleotide sequences of primers used for RT-qPCR, gene cloning, and genotyping.

